# An *in vitro* experimental pipeline to characterize the binding specificity of SARS-CoV-2 neutralizing antibodies

**DOI:** 10.1101/2023.04.20.537738

**Authors:** Kristina E. Atanasoff, Luca Brambilla, Daniel C. Adelsberg, Shreyas Kowdle, Christian S. Stevens, Chuan-Tien Hung, Yanwen Fu, Reyna Lim, Linh Tran, Robert Allen, J. Andrew Duty, Goran Bajic, Benhur Lee, Domenico Tortorella

## Abstract

The coronavirus disease 2019 (COVID-19) pandemic caused by the severe acute respiratory syndrome-coronavirus-2 (SARS-CoV-2) has led to over 760 million cases and >6.8 million deaths worldwide. We developed a panel of human neutralizing monoclonal antibodies (mAbs) targeting the SARS-CoV-2 Spike protein using Harbour H2L2 transgenic mice immunized with Spike receptor binding domain (RBD) (1). Representative antibodies from genetically-distinct families were evaluated for inhibition of replication-competent VSV expressing SARS-CoV-2 Spike (rcVSV-S) in place of VSV-G. One mAb (denoted FG-10A3) inhibited infection of all rcVSV-S variants; its therapeutically-modified version, STI-9167, inhibited infection of all tested SARS-CoV-2 variants, including Omicron BA.1 and BA.2, and limited virus proliferation *in vivo* (1). To characterize the binding specificity and epitope of FG-10A3, we generated mAb-resistant rcVSV-S virions and performed structural analysis of the antibody/antigen complex using cryo-EM. FG-10A3/STI-9167 is a Class 1 antibody that prevents Spike-ACE2 binding by engaging a region within the Spike receptor binding motif (RBM). Sequencing of mAb-resistant rcVSV-S virions identified F486 as a critical residue for mAb neutralization, with structural analysis revealing that both the variable heavy and light chains of STI-9167 bound the disulfide-stabilized 470-490 loop at the Spike RBD tip. Interestingly, substitutions at position 486 were later observed in emerging variants of concern BA.2.75.2 and XBB. This work provides a predictive modeling strategy to define the neutralizing capacity and limitations of mAb therapeutics against emerging SARS-CoV-2 variants.

**Importance:** The COVID-19 pandemic remains a significant public health concern for the global population; development and characterization of therapeutics, especially ones that are broadly effective, will continue to be essential as SARS-CoV-2 variants emerge. Neutralizing monoclonal antibodies remain an effective therapeutic strategy to prevent virus infection and spread with the caveat that they interact with the circulating variants. The epitope and binding specificity of a broadly neutralizing anti-SARS-CoV-2 Spike RBD antibody clone against many SARS-CoV-2 VOC was characterized by generating antibody-resistant virions coupled with cryo-EM structural analysis. This workflow can serve to predict the efficacy of antibody therapeutics against emerging variants and inform the design of therapeutics and vaccines.

## Introduction

The coronavirus disease 2019 (COVID-19) pandemic caused by severe acute respiratory syndrome-coronavirus-2 (SARS-CoV-2) is an unprecedented global public health challenge, with over 760 million cases and >6.8 million deaths worldwide as of March 2023 (2). In the last 18 years, bat coronaviruses SARS-CoV (2003), Middle East respiratory syndrome coronavirus (MERS-CoV) (2012), and SARS-CoV-2 (2019) have jumped from animal reservoirs to cause significant human outbreaks. The entry of SARS-CoV1 and SARS-CoV-2, as well as many other bat CoVs, is mediated via engagement of the envelope spike glycoprotein (denoted “Spike” or S) with the human angiotensin-converting enzyme 2 (ACE2) cell surface protein followed by activation and cleavage with transmembrane protease, serine 2 (TMPRSS2) (3–5). SARS-CoV-2 Spike is a type I membrane glycoprotein comprised of an S1 and S2 domain that binds to ACE2 via the receptor binding domain (RBD) contained within S1. Human monoclonal antibodies (mAbs) directed to the RBD of SARS-CoV-2 demonstrated efficacy in limiting disease symptoms and hospital stay as long as the mAbs recognized the circulating variants (6–11). However, due to the rise of new variants of concern (VoCs), the FDA’s Emergency Use Authorization of mAb therapeutics targeting Spike (e.g. LYCoV1404 (bebtelovimab) and tixagevimab/cilgavimab (Evusheld)) have been revoked due to demonstrated limited efficacy *in vitro* (12). Despite these findings, broadly neutralizing antibodies remain an effective therapeutic strategy to prevent virus infection and spread.

SARS-CoV-2 neutralizing monoclonal antibodies (nmAbs) target multiple regions of Spike to block virus binding to ACE2 or subsequent fusion with the host cell membrane. Such nmAbs may target either the S1 or S2 domain of Spike, with the most effective being those against the RBD within the S1 domain (13). NmAbs can be classified as binding to S1 (recognizing S1’s RBD or N-terminal domain) or S2 (recognizing the stem helix (SH) or fusion peptide (FP)). The anti-S1 nmAbs that interact with Spike RBD can be further subdivided into four classes: Class 1 and Class 2 nmAbs, which both overlap with the receptor-binding motif (RBM) and are distinguished by their ability to recognize RBD in exclusively the ‘up’ conformation of Spike (Class 1) or in both ‘up’ and ‘down’ conformations (Class 2); Class 3 nmAbs, which interact with the RBD outside of the RBM independent of Spike conformation; and Class 4 nmAbs, which recognize conserved regions within the RBD that do not directly block ACE2 binding. These nmAbs represent a large panel of biologicals whose neutralization capacity may diminish as VoCs arise. However, nmAbs that target conserved regions of Spike essential for virus fitness would likely maintain their therapeutic efficacy against emerging VoCs.

We recently generated a panel of nmAbs that inhibit SARS-CoV-2-mediated entry and infection. These nmAbs were elicited from Harbour H2L2® mice, which are transgenetically modified to encode human immunoglobulin variable regions and rodent constant regions; thus, immunization of these mice elicits a humoral response comprised of human nmAbs (14). From the panel of the nmAbs generated from Harbour H2L2 mice immunized with SARS-CoV-2 Spike (WA-1) RBD protein, one neutralizing monoclonal antibody, designated Family G (FG)-10A3 or its therapeutically-modified version STI-9167, had broad-spectrum neutralizing activity against currently circulating SARS-CoV-2 VoCs, including Omicron (BA.1 and BA.2), in both pseudotyped and live virus neutralization assays (1). To define the epitope of FG-10A3/STI-9167, we generated FG-10A3 mAb-resistant virions by utilizing replication-competent VSV expressing the SARS-CoV-2 WT-D614G (WA-1) Spike protein, and determined the structure of STI-9167:SARS-CoV-2 Spike (WA-1) complexes. Our study identified the phenylalanine 486 residue on the Spike protein critical for FG-10A3/STI-9167’s epitope recognition enabling the nmAb’s broad neutralizing activity against SARS-CoV-2 VOCs, while establishing a robust *in vitro* experimental strategy to define the binding specificity of mAb therapeutics and the molecular basis of the nmAb’s interaction with Spike.

## Material and Methods

### Cells and antibodies

Vero E6 (ATCC #CCL-81) and Vero angiotensin-converting enzyme 2 (ACE2) and transmembrane protease, serine 2 (TMPRSS2) expressing (Vero-ACE2/ TMPRSS2) cells (15) were maintained in Dulbecco’s modified Eagle’s medium (DMEM, Corning, 10-013-CV) supplemented with 10% heat-inactivated fetal bovine serum (FBS, ThermoFisher Scientific), 1 mM HEPES (Corning, 25-060-CI), 100 U/mL penicillin, and 100 g/mL streptomycin (100X Pen/Strep, Corning, 30-002-CI). Cell lines were cultured at 37°C with 5% CO_2_. Human monoclonal anti-SARS-CoV-2 Spike antibodies (mAbs) were generated, sequenced, and humanized, as described previously (1). The antibodies were purified from the supernatant of 293ExpiF cells transfected with the respective heavy and light chains of each clone. The FG-10A3 clone (Heavy chain CDR3: QVQLVESGGGVVQPGRSLRLSCAASGFTFSSYGMNWVRQAPGKGLEWVAIIWY<u>D</u>GNN<u>T</u>YYV DSVKGRFTISRDNSKNTLYLQMNSLRAEDTAVYYCARKD<u>G</u>SKTYYGYYFDYWGQGTLVTVSS; Light chain CDR3: DIQMTQSPSSLSASVGDRVTITCRASQSIHSFLNWYQQKPGKPPNLLIYAASSLQSGLPSRFSG SGSGTDFTLTISSLQPEDFATYYCQQSYITPPTFG<u>H</u>GTKVEIK) was modified with a LALA sequence to prevent ADE activity to generate the nmAb STI-9167 (STI-9167 (10A3YQYK) HC CDR123 AA sequence by kabat: QVQLVESGGGVVQPGRSLRLSCAASGFTFSSYGMNWVRQAPGKGLEWVAIIWY<u>Y</u>GNN<u>K</u>YYV DSVKGRFTISRDNSKNTLYLQMNSLRAEDTAVYYCARKD<u>Y</u>SKTYYGYYFDYWGQGTLVTVSS STI-9167 (10A3YQYK LC CDR123 AA sequence by kabat: DIQMTQSPSSLSASVGDRVTITCRASQSIHSFLNWYQQKPGKPPNLLIYAASSLQSGLPSRFSG SGSGTDFTLTISSLQPEDFATYYCQQSYITPPTFG<u>Q</u>GTKVEIK). The underlined residues represent modifications between the aa sequences.

### Sequencing of viral RNA

Viral RNA was extracted using the QIAamp Viral RNA Mini Kit (Qiagen). The Spike region of VSV-S virions was amplified via the SuperScript^TM^ IV One-Step RT-PCR System (Thermo Fisher Scientific) and purified using the SPRIselect - PCR Purification and Cleanup Kit (Beckman Coulter Life Sciences). Libraries were prepared using the Illumina DNA Prep Tagmentation Kit and barcoded using the Nextera DNA CD Indexes (Illumina). 150 bp paired-end sequencing was performed on an Illumina iSeq 100.

### Sequencing analysis

For bioinformatic analysis of raw sequencing data, an in-house analysis pipeline was used to process raw FASTQ files and identify all variants called at a threshold of 1%. Variants shown meet a threshold of 10% of all reads in any one sample at any point during passaging, result in an amino acid change, and are found at positions with a read depth of at least 5000 reads. First, reads were processed and mapped using SAMtools (16) and BWA-MEM (17) against SARS-CoV-2-Spike (WA-1) (18). Bcftools mpileup (17) and bedtools genomecov (19) were then used to identify all variants and their relative frequency across samples. Further analysis was performed in R using the tidyR (20) and Biostrings (21) packages.

### Generation of VSV-eGFP-CoV-2 Spike (Δ21 aa), point mutants and helper plasmids

We cloned the genomic sequences of all VSV-S pseudoviruses and helper plasmids into expression vectors as previously described (18). Briefly, we replaced the VSV-G open reading frame of a pEMC vector expressing VSV-eGFP with SARS-CoV2-S WT-D614G (WA-1) or specific variants B.135 (Beta), B.617.2 (Delta), and B.1.1.529 (Omicron BA.1), all of which were truncated and thus lacking the final 21 amino acids (22). The sequences of VSV-S(WA-1) and VSV-S(Beta) are available at Genebank (Accession Numbers: MW816497 and MW816499). We generated point mutants at the F486 residue by primer-mediated site-directed mutagenesis of the parental (WA-1) Spike. Forward and reverse primers were designed to mutate phenylalanine at position 486 to serine (F486S), leucine (F486L), or valine (F486V), and generate a 20bp overlap between fragments. Point-mutant Spike fragments were cloned into the MluI_PacI digested VSV-eGFP backbone via InFusion seamless cloning (TakaraBio). The sequences of VSV N, P, M, L were cloned into pCI vectors to generate helper plasmids for viral rescue. These accessory plasmids were a generous gift from Dr. Benjamin tenOever (NYU School of Medicine).

### VSV-eGFP-CoV-2 rescue from cDNA

For all VSV pseudoviruses, 3 x 10^5^ BHK-ACE2 cells per well were seeded onto 6-well plates. The next day, 2000 ng of pEMC-VSV-eGFP-CoV-2 Spike, 2500 ng of pCAGGS-T7opt (23), 850 ng of pCI-VSV-N, 400 ng of pCI-VSV-P, and 100 ng of pCI-VSV-L were mixed with 5.5 µL of Plus reagent and 8.9 µL of Lipofectamine LTX (Invitrogen) in 200 µL of OptiMEM medium (Gibco). Thirty minutes later, the transfection mixture was added dropwise onto plated BHK-ACE2 cells. Medium was replaced 24h post-transfection with DMEM supplemented with 10% FBS, and cells were monitored daily for fluorescence with a Celigo imaging cytometer (Nexcelom Bioscience). 3 to 5 days post-transfection, cells exhibited extensive GFP-positive syncytia and cell-free viral spread to previously uninfected cells. Supernatant containing pseudovirus was collected, clarified by centrifugation for 5 minutes at 400g, and amplified on Vero-CCL81-TMPRSS2 cells (15). The replication-competent (rc) VSV-S virions were titered on Vero-ACE2/TMPRSS2 cells based on number of GFP positive cells/ml.

### VSV-S neutralization assays

Vero-ACE2/TMPRSS2 cells (1×10^4^/well) were plated in a 96-well plate and placed at 37°C/5% CO^2^ overnight (∼18hrs). The cells were infected with rcVSV-S virions (MOI=0.2 in 50 µL DMEM) pre-incubated with anti-Spike mAbs (0, 1.4, 4.1, 12.3, 37, 111, 333, 1000 ng/mL at 2X concentrations in 50 µL DMEM) at 4°C for 1 hr. The total 100 µL of virus mixed with mAbs was then added to cells, followed by incubation for ∼18hpi at 37°C. Cells were fixed in 4% paraformaldehyde (PFA), permeabilized (0.3% Triton X-100 (ThermoFisher, HFH10) in PBS), and then stained with Hoechst reagent (0.01 µg/mL in PBS) to quantify total cells. Virus neutralization was quantified using a plate-based imaging Celigo cytometer (Nexelcom Bioscience, Version 4.1.3.0) based on the number of GFP-positive cells as well as the total number of cells per well. Relative percent infection was determined using the number of GFP+ cells in wells infected, with VSV-S alone as 100% infection.

### Generation of mAb-resistant VSV-S(WA-1) virions

Vero E6 cells (2.5×10^4^/well) were plated in a 24-well plate and placed at 37°C/5% CO_2_ overnight (∼18hrs). The cells were infected with rcVSV-S(WA-1) (MOI=0.1 in 500 µL DMEM) pre-incubated with anti-Spike nmAb FG-10A3 (0, 0.15, 0.46, 1.4, 4.1, 12.3, 37, 111, 333, 1000, 3000 ng/mL final concentrations in 1 ml DMEM) at 4°C for 1hr. The total 1 mL of virus mixed with mAbs was added to cells, followed by incubation at 37°C. At 72 hours post-infection, the percentage of GFP-positive cells was determined using a Celigo cytometer, and the supernatant from wells with >50% of infection (50 µL) was selected for incubation with FG-10A3 (0-3000 ng/ml). Passaged virus was defined as “resistant” to FG-10A3 when the virus from the >50% infection well required >600 ng/ml of FG-10A3 (100*EC_50_ (6 ng/ml)) to achieve >50% infection. Once resistance was elicited, virus was passaged twice more with FG-10A3 (0-3000 ng/ml) to verify that viral resistance could be maintained yielding a total of 8 viral passages. Stocks of these 8 passages were grown and amplified in the absence of nmAb, then titered on Vero-ACE2/TMPRSS2 cells prior to assessment of nmAb-mediated neutralization. To control for amino acids changes that arise through rcVSV-S passaging, rcVSV-S WA-1 incubated with 1000 ng/ml of an isotype control was collected over 8 passages.

### ACE2/antibody competition assay

Purified monoclonal antibodies (0.1 and 1μg/ml) were pre-incubated with Wuhan Spike RBD-mG2a Fc fusion protein (0.25mg/ml) for 30 minutes on ice in FACS Buffer (1xPBS, 0.5% BSA, 2mM EDTA) followed by addition to HEK-293 cells expressing human ACE2 for 30 minutes on ice. Cells were washed 2x with FACS Buffer and resuspended in FACS Buffer containing goat anti-mouse IgG-APC secondary (1:1000). Cells were incubated for 30 minutes on ice, washed 2x with FACS Buffer and resuspended in FACS buffer for analysis on a Intellicyte HFTC (in duplicate). Mean fluorescent intensities was determined and normalized using RBD as 100% binding. Secondary alone was included as a negative control.

### K_D_ determination

Biolayer interferometry assays were performed using the Octet RED 96 instrument (SartoriusAG) to determine the association rates (k_on_), dissociation rates (k_dis_), and affinity (K_D_) for the antibody. Purified Spike receptor binding domain (RBD)-mouse Fc fusion protein onto an anti-mouse Fc IgG capture (AMC) biosensors using constant 5μg/mL concentration for 10 min at 20°C. To determine the kon, the sensors were exposed to the human antibodies starting at a concentration of 1 ug/mL (2-fold dilutions (1-0.0078 ug/mL in PBS) for 3 min. To determine the K_D_s, dissociation was measured over the course of 3 min while the sensors were in PBS buffer. K_D_ values were calculated based on the as a ratio of k_dis_/k_on_. A binding model of 1:1 resulted in the best fit for each antibody and the resulting R2 values are given for each antibody in Table 1.

**Table 1.** Antibody diversity and SARS-CoV-2 Spike RBD binding affinity for nmAbs.

|  |  |  |  |  |
| --- | --- | --- | --- | --- |
| **Antibody** | **CDR3 (H)** | **VDJ (Heavy)** | **VJ (Kappa)** | **K<sub>D</sub> (M)** |
| FA-10D6 | 12 aa | VH3-53/DH6-19/JH3 | VK1-9/JK2 | 5.74 E-11 |
| FB-1D10 | 19 aa | VH1-8/DH3-10/JH6 | VK3-20/JK5 | 5.63 E-11 |
| FD-2C1 | 14 aa | VH3-49/DH6-13/JH4 | VK3-15/JK1 | 3.71 E-10 |
| FE-14G5 | 16 aa | VH4-4/DH3-19/JH4 | VK3-15/JK2 | 1.12 E-10 |
| FG-10A3 | 16 aa | VH3-33/DH2-21/JH4 | VK1-39/JK1 | 1.25 E-10 |

### Cryo-EM sample preparation and data collection

SARS-CoV-2 HexaPro Spike (24) was incubated with STI-9167 Fab at 2.5 mg/mL and a molar ratio of 1.5:1 Fab:Spike for 20 minutes at 4°C. Immediately before grid preparation, fluorinated octyl maltoside was added to the pre-formed complex at 0.02% w/v final concentration. Then, 3 µL aliquots were applied to UltrAuFoil gold R0.6/1 grids and subsequently blotted for 6 seconds at blot force 1, then plunge-frozen in liquid ethane using an FEI Vitrobot Mark IV. Grids were imaged on a Titan Krios microscope operated at 300 kV and equipped with a Gatan K3 Summit direct detector. 20,590 movies were collected in counting mode at 16e−/pix/s at a magnification of 105,000, corresponding to a calibrated pixel size of 0.826 Å. Defocus values were from -0.5 to -1.5 μm.

### Cryo-EM data processing

Movies were aligned and dose weighted using MotionCorr2 (25) Contrast transfer function estimation was done in cryoSPARC v3.3.1 using Patch CTF, and particles were picked with cryoSPARC’s blob picker. The picked particles were extracted with a box size of 512 pixels, with 4x binning and subjected to a 2D classification. Selected particles were then subjected to a second round of 2D classification. An initial model was generated on the 1,410,814 selected particles at 6 Å/pixel with 4 classes. The best class, containing 558,457 particles, was selected for further processing. After one round of non-uniform refinement, without imposed symmetry, the particles were subjected to 3D classification with 5 classes. Of these, the best 3 classes, containing 321,918 particles in total, were combined, re-extracted without binning with a box size of 512 pixels, and selected for further rounds of non-uniform refinement with local CTF refinement, yielding the final global map at a nominal resolution of 2.53 Å. The protomer with the best Fab volume was subjected to local refinement with a soft mask extended by 6 pixels and padded by 12 pixels encompassing the RBD and Fab. A second round of local refinement was performed with a soft mask encompassing the RBD and variable domains of the Fab. This yielded the final local map at 3.16 Å resolution. The two half-maps from the global or local refinement were used for sharpening in DeepEMhancer (26). The reported resolutions are based on the gold-standard Fourier shell correlation of 0.143 criterion.

### Model building and refinement

RBD from PDB (ID: 6M0J) and AlphaFold2-predicted Fab variable domains were fit into the focus refined maps using UCSF ChimeraX (27) and then manually built using COOT (28). N-linked glycans were built manually in COOT using the glyco extension and their stereochemistry and fit to the map validated with Privateer (29). The model was then refined in Phenix (30) using real-space refinement and validated with MolProbity (31).

### Statistics and reproducibility

All statistical tests were performed using GraphPad Prism 9 software (La Jolla, CA). The half-maximal effective concentration (EC_50_) values for each anti-Spike mAb were calculated using 3-parameter non-linear regression analysis after the mAb concentrations were transformed to log scale. Significance and adjusted p-values for mAbs’ inhibitory effects on infection were calculated via 2-way ANOVA statistical tests, where the mean relative percent infection of each experimental condition was compared to the mean relative percent infection of cells treated with virus alone. For all figures, error bars represent standard deviation of the mean. Sample size and replicates for each experiment are indicated in the figure legends. Technical replicates were prepared in parallel within one experiment, and experimental replicates were performed on separate days.

## Results

### Human anti-RBD neutralizing mAbs block infection of rcVSV-S variants

A panel of human neutralizing monoclonal antibodies (nmAbs) against SARS-CoV-2 was identified from Harbour H2L2® mice immunized with a receptor binding domain (RBD)-mouse Fc fusion protein (1). Hybridoma clones demonstrating: 1) binding to 293ExpiF cells transfected with SARS-CoV-2 spike cDNA with 5 fold greater than background; 2) RBD-binding as determined by ELISA; and 3) rcVSV-S(WA-1) neutralization of >50% were selected for nucleotide sequence analysis to identify unique clones (1). The neutralizing clones were classified into antibodies families based on amino acid sequence, V(D)J gene usage, and complementarity-determining region 3 (CDR3) junction identity. The CDR3 lengths ranged from 12-19 aa in length, with subnanomolar measured affinities (K_D_s) against SARS-CoV-2 Spike RBD ranging from 0.057-0.13 nM (**Table 1**). Collectively, this elicited a panel of genetically distinct human nmAbs targeting the SARS-CoV-2 Spike RBD.

We selected clones of Family A, B, D, E, and G antibodies based on their robust neutralization activity to evaluate broad neutralization based on neutralization activities against the SARS-CoV-2 (WA-1) (1). To evaluate the neutralizing capacity of the anti-RBD antibodies, we generated rcVSV-S virions expressing the WT-D614G (WA-1), B.135 (Beta), B.617.2 (Delta), and BA.1 (Omicron) Spike variants as a proxy to examine neutralization under BSL2 conditions. The rcVSV-S virions were preincubated with representative clones of Family A clone 10D6 (FA-10D6), B clone FB-1D10 (FB-1D10), D clone 2C1 (FD-2C1), E clone 14G5 (FE-14G5), and G clone 14G5 (FE-14G5) nmAbs (0-1 µg/mL) prior to infection of Vero-ACE2-TMPRSS2 cells and analyzed for GFP fluorescence by a Celigo cytometer (**Figure 1**). The relative percent infection was determined using pretreatment with an isotype mAb control as 100% infection. The results revealed diverse neutralization profiles among the antibodies. FA-10D6 and FD-2C1 inhibited infection of rcVSV-S WA-1 and Delta suggesting these mAbs recognize RBD regions conserved between the WA-1 and Delta Spike proteins (**Figure 1A and C**). On the other hand, FB-1D10 neutralized rcVSV-S WA-1 and Beta, while FE-14G5 limited infection mediated by rcVSV-S WA-1 and Omicron BA.1, respectively (**Figure 1B and D**). Strikingly, FG-10A3 demonstrated broad neutralization of infection by rcVSV-S WA-1, Beta, Delta, and Omicron (**Figure 1E**) with EC_50_ values ranging from 6.2-11 ng/mL (**Table 2**) implying the antibody recognizes an epitope conserved among all variants. The ability of FG-10A3 to broadly neutralize rcVSV-S variants is consistent with the findings that an engineered version of FG-10A3 denoted STI-9167 neutralized the corresponding SARS-CoV-2 VoCs (1). As such, identifying FG-10A3’s epitope will define a RBD region conserved across different variants that can be targeted by nmAb-based therapeutics.

**Figure 1.**
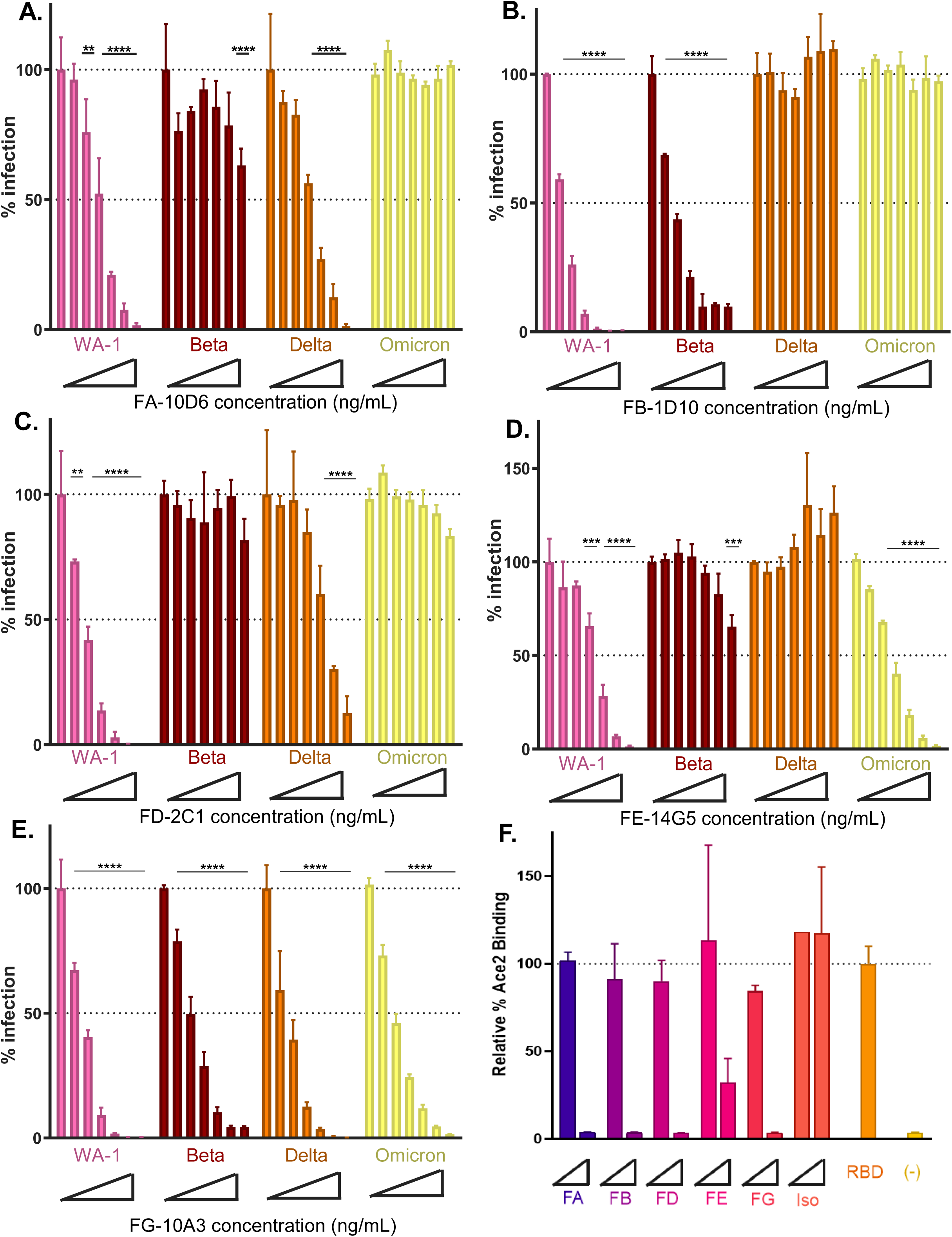
Analysis of anti-SARS-CoV-2 nmAbs against VSV-S variants. FA-10D6 (**A**), FB-1D10 (**B**), FD-2C1 (**C**), FE-14G5 (**D**), and FG-10A3 (**E**) were evaluated with a neutralization assay against rcVSV-S WA-1 (WA-1), rcVSV-S variant B.135 (β, Beta), rcVSV-S variant B.617.2 (δ, Delta), and rcVSV-S variant B.1.1.529 (ο, Omicron). rcVSV-S virions (MOI=0.2) were preincubated with increasing concentrations of nmAbs (0-1 µg/mL) followed by infection of Vero-ACE2/TMPRSS2 cells and analyzed for infection at 18hpi. Infection was assessed by the number of GFP-positive cells and 100% infection was determined from untreated rcVSV-S, respectively. Error bars represent standard deviation from the mean of three samples. Statistical significance is denoted as: **, *p*<0.01; ***, *p*<0.001; ****, *p*<0.0001. (**F**) Recombinant SARS-CoV-2(WA-1) RBD-Fc was preincubated with 0.1 and 1 µg/mL of FA-10D6, FB-1D10, FD-2C1, FE-14GE, or FG-10A3 and then added to HEK-293 cells expressing hACE2 followed by flow cytometry analysis. The relative percent ACE2 binding was determined using untreated RBD-Fc binding to ACE2 expressing cells as 100%. Error bars represent standard deviation from the mean of duplicate samples.

**Table 2.** Half-maximal inhibitory concentrations (IC50) for anti-RBD nmAbs against rcVSV-S variants.

|  |  |  |  |  |  |  |  |
| --- | --- | --- | --- | --- | --- | --- | --- |
|  | **rcVSV-S variant (IC<sub>50</sub> (ng/mL))** |  |  |  |  |  |  |
| **nmAb** | **WA-1** | **Beta** | **Delta** | **Omicron** | **WA-1 + F486S** | **WA-1 + F486L** | **WA-1 + F486V** |
| FA-10D6 | 33.5 | n/a | 54.7 | n/a | 101.8 | 13.5 | 24.4 |
| FB-1D10 | 2.19 | 6.42 | n/a | n/a | n/a | n/a | n/a |
| FD-2C1 | 6.2 | n/a | 173.4 | n/a | 110.2 | 2.45 | 2.84 |
| FE-14G5 | 72.7 | n/a | n/a | 26.4 | 75.9 | 16.8 | 12.7 |
| FG-10A3 | 6.21 | 11 | 9.31 | 10.8 | n/a | 150.1 | 102.6 |

We next evaluated the inhibitory function of the anti-RBD nmAbs using an ACE2 binding assay. Recombinant SARS-CoV-2 (WA-1) RBD-Fc preincubated with FA-10D6, FB-1D10, FD-2C1, FE-14G5, FG-10A3, or non-RBD control antibody were added to HEK-293 cells expressing ACE2, followed by flow cytometry analysis for RBD-Fc binding to ACE2 (**Figure 1F**). The binding of RBD-Fc to ACE2-expressing cells yielded a strong fluorescence signal that was dramatically reduced upon preincubation with anti-RBD nmAbs. These data indicate that the nmAbs elicited against RBD limited SARS-CoV-2 infection by preventing Spike binding to ACE2.

### Generation of resistant rcVSV-S WA-1 virions against the anti-RBD nmAb FG-10A3

Based on the broad neutralization of FG-10A3 against SARS-CoV-2 VOCs, we propose that the antibody interacts with a conserved epitope among the spike VOCs. Thus, we planned to define the critical resides of the Spike protein that facilitate its interaction with FG-10A3. A strategy to identify the critical residues of a mAb’s protein target is to generate antibody-resistant virus strains, followed by genomic sequencing to identify candidate resistance-conferring amino acid changes (32). The amplification of rcVSV-S in the presence of the nmAb will provide a selective context for identification of antibody-resistant virus over several passages. To that end, we utilized rcVSV-S WA-1 to select for mutants resistant to FG-10A3 (**Figure 2**). Briefly, rcVSV-S WA-1 infected Vero-E6 cells were incubated with FG-10A3 (0-3 µg/mL) for three days followed by analysis of GFP positive cells using a Celigo imaging cytometer. Supernatant from the well with the highest nmAb concentration that revealed a >50% infection was selected for additional rounds of incubation with the range of FG-10A3 concentration (**Figure 2A**). Using this strategy of collecting and passaging virus from cells demonstrating ∼50% infection, resistant rcVSV-S WA-1 was observed at passage 6, and we continued to collect virus from an additional two passages. To validate the generation of antibody-resistant rcVSV-S WA-1, we evaluated the neutralization capacity of FG-10A3 against virions collected from all passages (**Figure 2B**). FG-10A3 neutralized virus harvested from rcVSV-S WA-1 Passages 1-5 with EC_50_ values comparable to the parent rcVSV-S WA-1 strain. However, FG-10A3 was unable to inhibit infection of virus collected from Passages 6-8 implying that the rcVSV-S WA-1 virions were insensitive to FG-10A3. Thus, the findings support the generation of FG-10A3 resistance rcVSV-S WA-1 virions.

**Figure 2.**
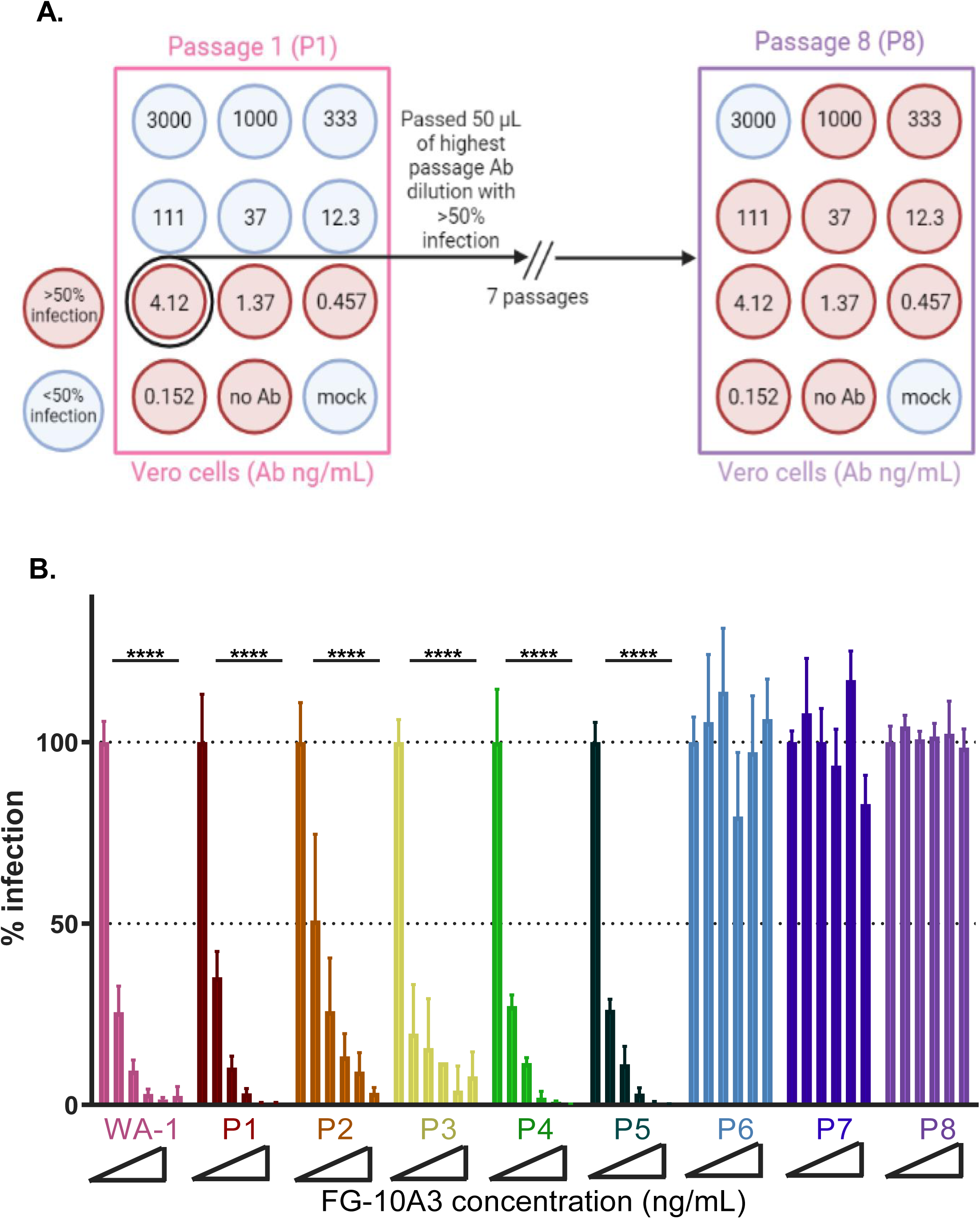
Generation of rcVSV-S WA-1 virions resistant to FG-10A3. (**A**) Schematic diagram illustrates the strategy to generate FG-10A3 resistant rcVSV-S WA-1 virions. Vero E6 cells infected with rcVSV-S WA-1 (MOI=0.1) following the incubation with FG-10A3 (0-3000 ng/mL) were assessed for GFP positive cells with the Celigo cytometer at 72hpi. Infection was assessed by the number of GFP-positive cells with untreated rcVSV-S WA-1 as 100% infection. Supernatant from wells with >50% infection treated with the highest concentration of FG-10A3 was considered Passage 1 (P1) and selected for subsequential incubation with FG-10A3 (0-3 µg/mL). Using this strategy, rcVSV-S WA-1 was passaged for a total of 8 passages, denoted P1-P8. (**B**) FG-10A3-incubated RcVSV-S from P1-P8 was assessed using an FG-10A3-based neutralization assay with 100% infection determined from untreated rcVSV-S WA-1. Error bars represent standard deviation from the mean of three samples. Statistical significance is denoted as: ****, *p*<0.0001.

### Identification of mutations within RBD responsible for the FG-10A3-resistant to rcVSV-S WA-1 virions

Viral RNA was extracted from Passage 1-8 of rcVSV-S WA-1 incubated with the isotype control or FG-10A3. The Spike region of rcVSV-S WA-1 virions was PCR amplified and sequenced at a total depth of 7.7 million reads across all samples, with 4.9 million aligning to SARS-CoV-2 Spike. Samples contained between 10.7k and 75.2k aligned reads, with a median of 17.3k. After variant calling, only those variants both resulting in an amino acid change and meeting a threshold of 10% of the total reads in any one sample are shown (**Figure 3**). Importantly, analysis of mutations in the RBD (aa. 319-541) identified changes at position F486 that were unique only to the FG-10A3 incubated rcVSV-S WA-1 virions compared to isotype control (**Figure 3A versus 3B**). A mutation from F to L arose at Passages 5 and 6 and then outcompeted by a change to S at Passage 6 which reached >99% at Passage 7 (**Figure 3B**). These findings suggest that F486 is a key residue for FG-10A3 neutralization of SARS-COV-2. Interestingly, rcVSV-S WA-1 passaged in the presence of isotype control antibody or FG-10A3 identified mutations in the S1 (aa. 1-685) and S2 (aa. 686-1273) domains of the spike protein (**Figure 3**). The spike H69R, Q183K, S248R, H655L, R685G, G769E, and R1185W mutations were observed during the passage of rcVSV-S WA-1 incubated with isotype antibody (**Figure 3A**), while H655L, S691G, and G769E were identified rcVSV-S WA-1 virions passaged with FG-10A3 (**Figure 3B**). These amino acid changes likely arose to enhance proliferation and dissemination of rcVSV-S WA-1 *in vitro* through likely impacting Spike binding and processing during infection and spread. Collectively, the experimental strategy of sequencing mAb-resistant virus strains generated *in vitro* can identify residues of the target viral protein that are critical for antibody binding.

**Figure 3.**
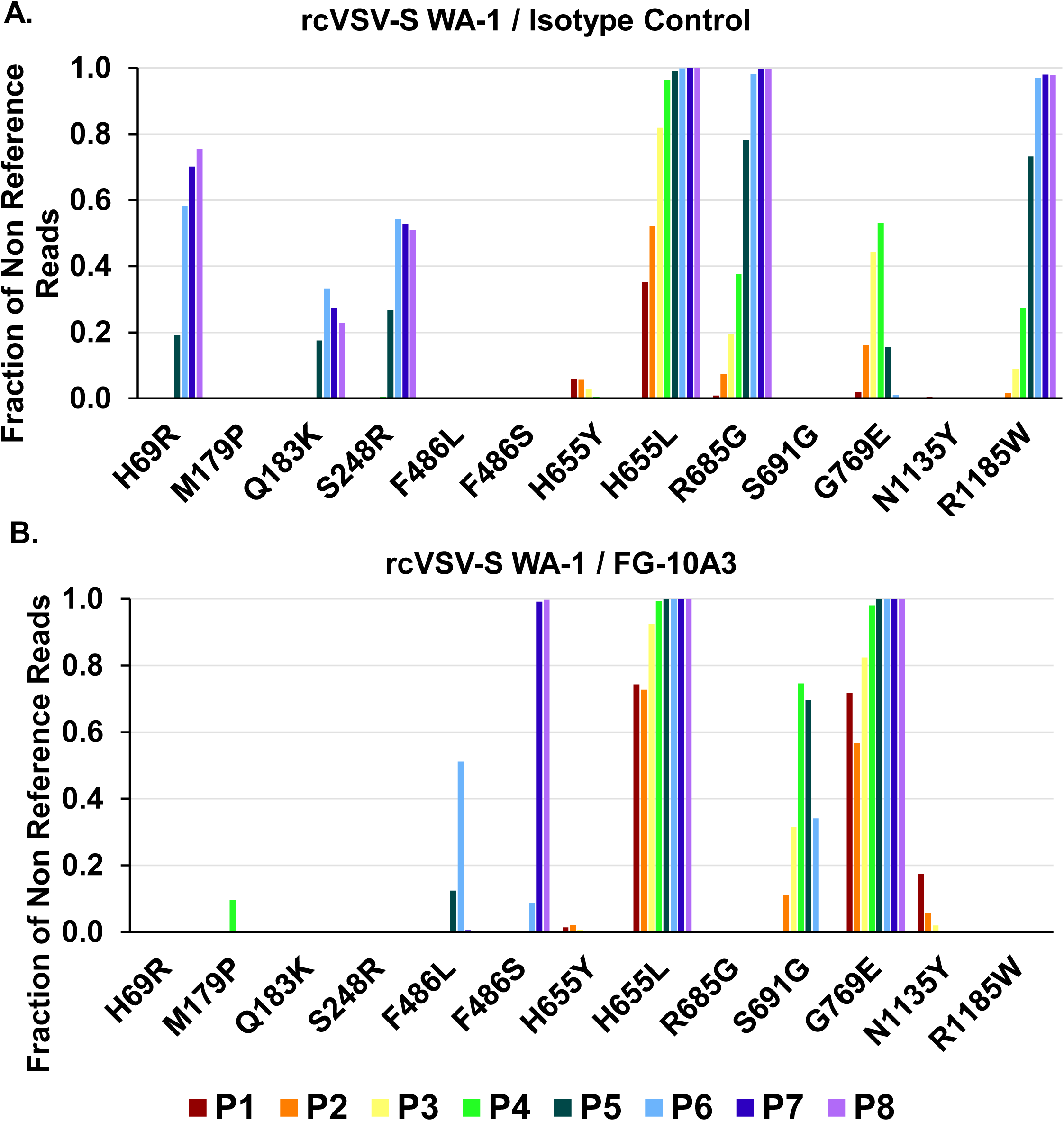
Identification of amino acid changes in rcVSV-S WA-1 virions resistant to FG-10A3. Viral RNA extracted from 8 sequential passages of rcVSV-S WA-1 (**A**) or rcVSV-S WA-1 incubated with FG-10A3 (**B**) was amplified via RT-PCR and subjected to Illumina sequencing and analysis of Spike mutational variants. The changes of amino acids at different residues of the spike protein are represented by the single letter amino acid code as a fraction of reads compared to non reference reads. The sequencing of rcVSV-S WA-1 from the Passage (P) 1-8 are shown in different colors.

### A polar residue at Spike position 486 dramatically reduces the neutralization capacity of FG-10A3

To validate the importance of F486 in FG-10A3-mediated neutralization, we performed neutralization assays using rcVSV-S WA-1 variants in which the F486 was modified to a leucine (rcVSV-S F486L), valine (rcVSV-S F486V), or serine (rcVSV-S F486S) (**Figure 4A**). RcVSV-S F486S was resistant to neutralization by FG-10A3 supporting the findings of our FG-10A3 resistant rcVSV-S WA-1 study (**Figures 2 ad 3**). Interestingly, neutralization of rcVSV-S F486L and rcVSV-S F486V by FG-10A3 was only somewhat reduced compared to the parental rcVSV-S WA-1 to EC_50_ values of 150.1 ng/mL and 102.6 ng/mL, respectively (**Table 2**). These results suggest that the substitution of a phenylalanine to nonpolar residues leucine or valine only slightly impacted nmAb binding to Spike while still permitting antibody-mediated virus neutralization. Hence, the conservation of the hydrophobic nature of residue 486 within the Spike protein would continue to be a target for FG-10A3.

**Figure 4.**
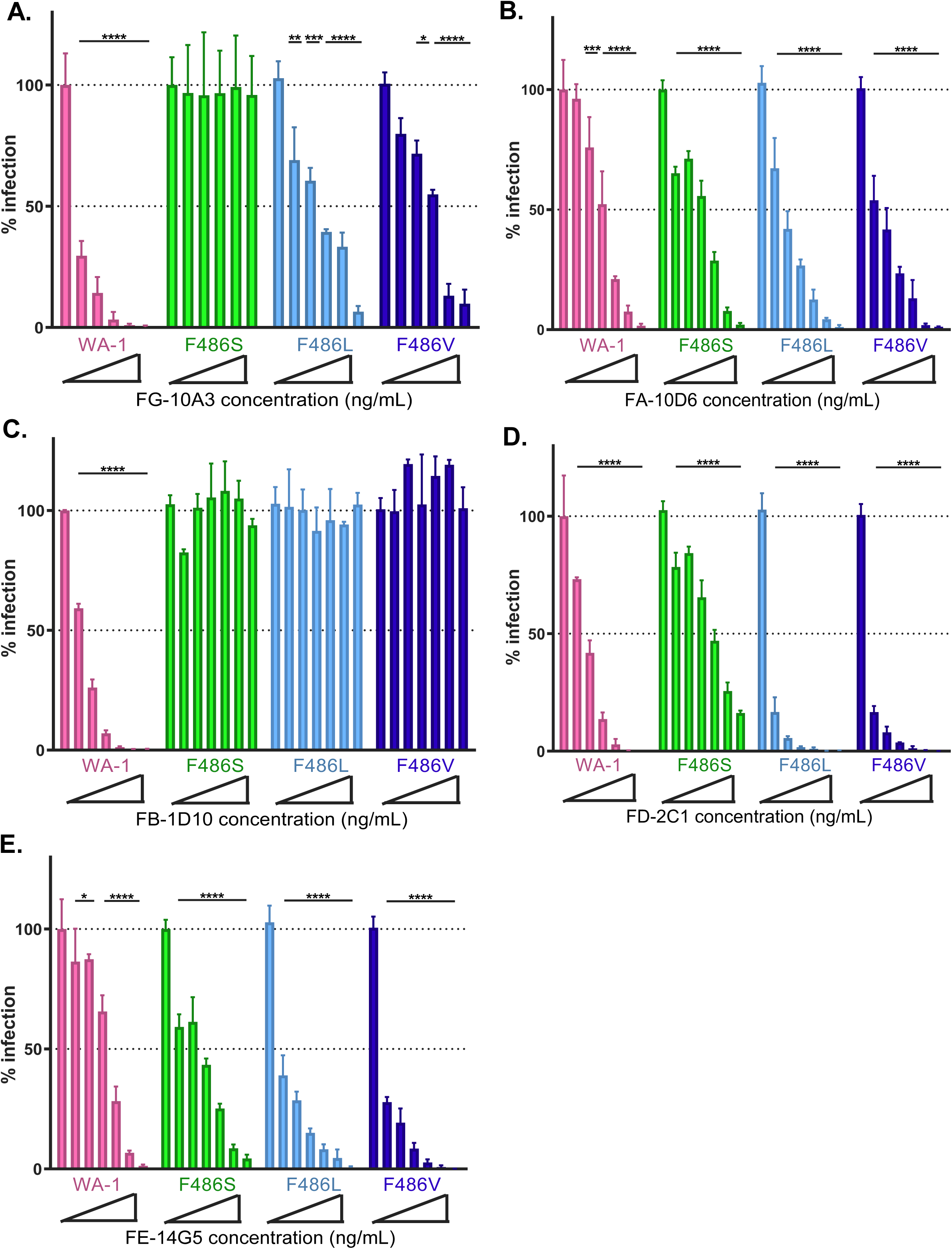
Analysis of anti-SARS-CoV-2 nmAbs against VSV-S F486 point mutants. FG-10A3 (**A**), FA-10D6 (**B**), FB-1D10 (**C**), FD-2C1 (**D**), and FE-14G5 (**E**) were evaluated with a neutralization assay against rcVSV-S WA-1 and rcVSV-S WA-1 with point mutants F486S, F486L, and F486V. Infection was assessed by the number of GFP-positive cells and 100% infection was determined from untreated rcVSV-S WA-1, respectively. Error bars represent standard deviation from the mean of three samples. Statistical significance is denoted as: *, *p*<0.05; **, *p*<0.01; ***, *p*<0.001; ****, *p*<0.0001.

Do mutations of Spike at position F486 impact the neutralization capacity of other anti-RBD nmAbs? To address this question, we evaluated the neutralization capacity of FA-10D6, FB-1D10, FD-2C1, and FE-14G5 against rcVSV-S WA-1 carrying F486 point mutants (**Figure 4B-F**). rcVSV-S WA-1, F486S, F486L, and F486V preincubated with anti-RBD nmAbs (0-1 µg/mL) prior to infection of Vero-ACE2-TMPRSS2 cells were analyzed for GFP fluorescence by a Celigo imaging cytometer, and relative percent infection was determined using a non-neutralizing isotype control as 100% infection. FA-10D6, FD-2C1, and FE-14G5 limited infection of all F486 point mutants, implying these nmAbs target a different epitope of the RBD than FG-10A3, and moreover that these point mutations at residue 486 do not induce a major conformational change within the Spike protein (**Table 2**). However, FB-1D10 was unable to neutralize any of the F486 mutants, suggesting that F486 is a critical residue for contact with this nmAb. Together, these data indicate that these RBD-targeting nmAbs have unique epitopes, and that a single amino acid change can significantly impact mAb neutralization activity.

### Cryo-EM structure of STI-9167 in complex with SARS-CoV-2 Spike

To define the epitope and the intermolecular interactions occurring at the antibody:antigen interface, we determined the structure of STI-9167 Fab (a therapeutically-modified version of FG-10A3; see (1)) in complex with SARS-CoV-2 Spike using single-particle cryo-EM (**Figure 5, Figure S1, Table S1**). The global structure indicated that STI-9167 binds to Spike along its three-fold axis and engages the RBD in the ‘up’ configuration (**Figure 5A**). We next performed local refinement of the STI-9167 Fab/RBD complex to resolve the amino acid side-chain contacts at the interface. The locally refined map, at 3.16 Å nominal resolution, indicated that the Fab contacts the disulfide-stabilized 470-490 loop with both its variable heavy and light chains (**Figure 5B-E**). The buried interaction surface area of 610 Å^2^ is relatively small compared to other anti-RBD nmAbs, such as bamlanivimab, which has an interaction surface area of 836 Å^2^.

**Figure 5.**
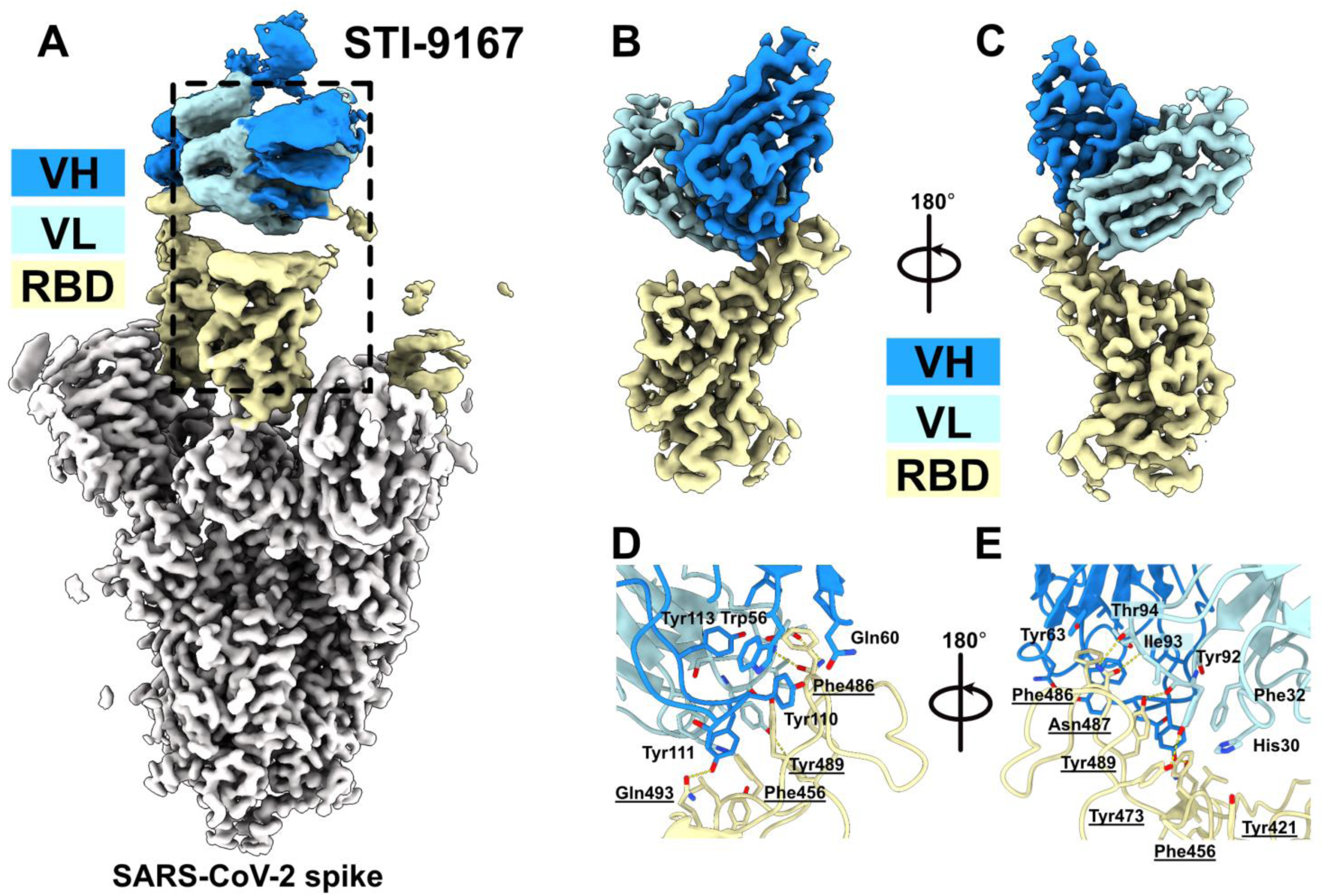
Structural analysis of anti-Spike RBD nmAb with SARS-CoV-2 Spike. Global (**A**) and local (**B, C**) cryo-EM reconstructions of the STI-9167 Fab in complex with SARS-CoV-2 Spike are shown with RBD colored in yellow, non-RBD regions in grey, and Fab variable heavy and light chain in bold and light blue, respectively. Two 180-degree rotated views (**D, E**) display details of the intermolecular interactions, with the most prominent interacting residues annotated. RBD-interacting residues are both numerically notated and underscored.

At the nexus of the interactions within the antibody:antigen interface is the RBD residue F486, which protrudes into a hydrophobic cleft formed by aromatic residues supplied predominantly by the HCDR2 and HCDR3 antibody loops. Indeed, the antibody residues W56, Y63, Y111 and Y113 wrap around the RBD F486, bolstering our findings with the rcVSV-S F486 point mutants. To wit, FG-10A3/STI-9167 can tolerate substitutions on RBD at position 486 so long as it is another hydrophobic residue; however, introduction of a polar residue results in the loss of hydrophobic interactions at this interface, explaining the total impairment of FG-10A3-mediated neutralization upon introduction of the F486S mutation.

In addition, most polar contacts within this interface involve the Fab light chain, which forms hydrogen bonds with RBD Y473, N487 and Y489. Interestingly, the majority of polar contacts involve the main chain carbonyls supplied by the mAb LCDR3 Y92, I93 and T94 residues (**Figure 5E**). In summary, the small binding interface, the main-chain contacts, and the high viral mutational cost at position 486 all contribute to the broadly reactive and potent neutralizing activity of nmAb FG-10A3/STI-9167 against SARS-CoV-2 VoCs.

## Discussion

A novel coronavirus was identified in December 2019 as the causative agent of the disease referred to as COVID-19 (33). In the intervening time since its isolation, the coronavirus SARS-CoV-2 has spread to virtually every part of the globe, while also dramatically expanding with regards to its catalogue of circulating “variant” strains, especially those containing mutational variation within viral proteins such as the Spike protein. Defining and evaluating such mutations is important for predicting and assessing the continued efficacy of existing antiviral therapeutics and vaccine efficacy. In this work, we have utilized two diverse approaches to identify the epitope of the broadly reactive anti-RBD nmAb, FG-10A3/STI-9167. A residue within the RBD that is critical for FG-10A3 neutralizing capacity was uncovered by experimentally generating FG-10A3 resistant rcVSV-S WA-1 virions (**Figures 2, 3, and 4**), in conjunction with cryo-EM structural analysis of STI-9167 in complex with the Spike protein (**Figure 5**). Together, these data support the importance of residue F486 within Spike as a contact residue with FG-10A3/STI-9167. The experimental pipeline utilized to identify the epitope of a broadly neutralizing anti-SARS-CoV-2 mAb highlights a powerful *in vitro* strategy to define the therapeutic potential of biologics for current and future viral variants.

The cryo-EM structural analysis of STI-9167 with Spike revealed contact residues at the disulfide-stabilized 470-490 loop in the Spike RBD (**Figure 5**). The hydrophobic interactions between F486 and numerous aromatic residues of the antibody heavy chain are at the heart of the antibody:antigen interface, with additional polar contacts supplied by the antibody light chain. The STI-9167:Spike complex highlights the unique interactions that allow for the antibody to broadly neutralize numerous VoCs. The importance of the F486 residue for interactions with FG-10A3/STI-9167 was supported by the generation of rcVSV-S resistant to FG-10A3, in which complete resistance was acquired through the F486S mutation and further validated by a rcVSV-S neutralization assay (**Figure 4**).

At the time of these studies, all SARS-CoV-2 VoCs were sensitive to FG-10A3/STI-9167 due to an overall lack of mutational changes within Spike RBD residues that make contact with the antibody. However, some SARS-CoV-2 subvariant lineages, namely BA.2.75.2 and XBB.1.5, include the mutation F486S or F486P, respectively, (**Figure 6**) suggesting that these VOCs would be insensitive to FG-10A3/STI-9167-mediated neutralization. Indeed, substitutions at position 486, including the F486S substitution, confer significant antibody evasion capability in both circulating SARS-CoV-2 variants (34) and laboratory-generated viral escape mutants solicited under nmAb-directed selective pressure (35). Importantly, the F486S mutation may incur a fitness “penalty” for the virus, in the form of impaired binding to its ACE2 cellular receptor (36). The structure of SARS-CoV-2 Spike RBD in association with ACE2 supports the idea that the presence of a large, hydrophobic residue at Spike position 486 facilitates efficient receptor binding via engagement with a hydrophobic “pocket” made by ACE2 residues L79, M82, and Y83 (37). This perhaps contributes toward explaining why, despite F486S’s significant fitness advantage with regards to antibody evasion, this mutation has only been found in ∼0.1-2% of circulating variants worldwide (38), whereas a substitution such as F486V, which maintains the favorable RBD:ACE2 hydrophobic interactions, has been uncovered in approximately 15-40% of circulating variants (39). Thus, consideration of the totality of the evolutionary “landscape” is critical when attempting to assess the viability and relevance of SARS-CoV-2 mutational variations in the context of *in vivo* pathogenesis. By contrast, the relatively permissive fitness environment established by our *in vitro* rcVSV reporter system permits a broader, more enhanced identification of potential limitations to a therapeutic’s efficacy that may lie outside of the rigors of selection imposed by the pre-existing evolutionary environment.

**Figure 6.**
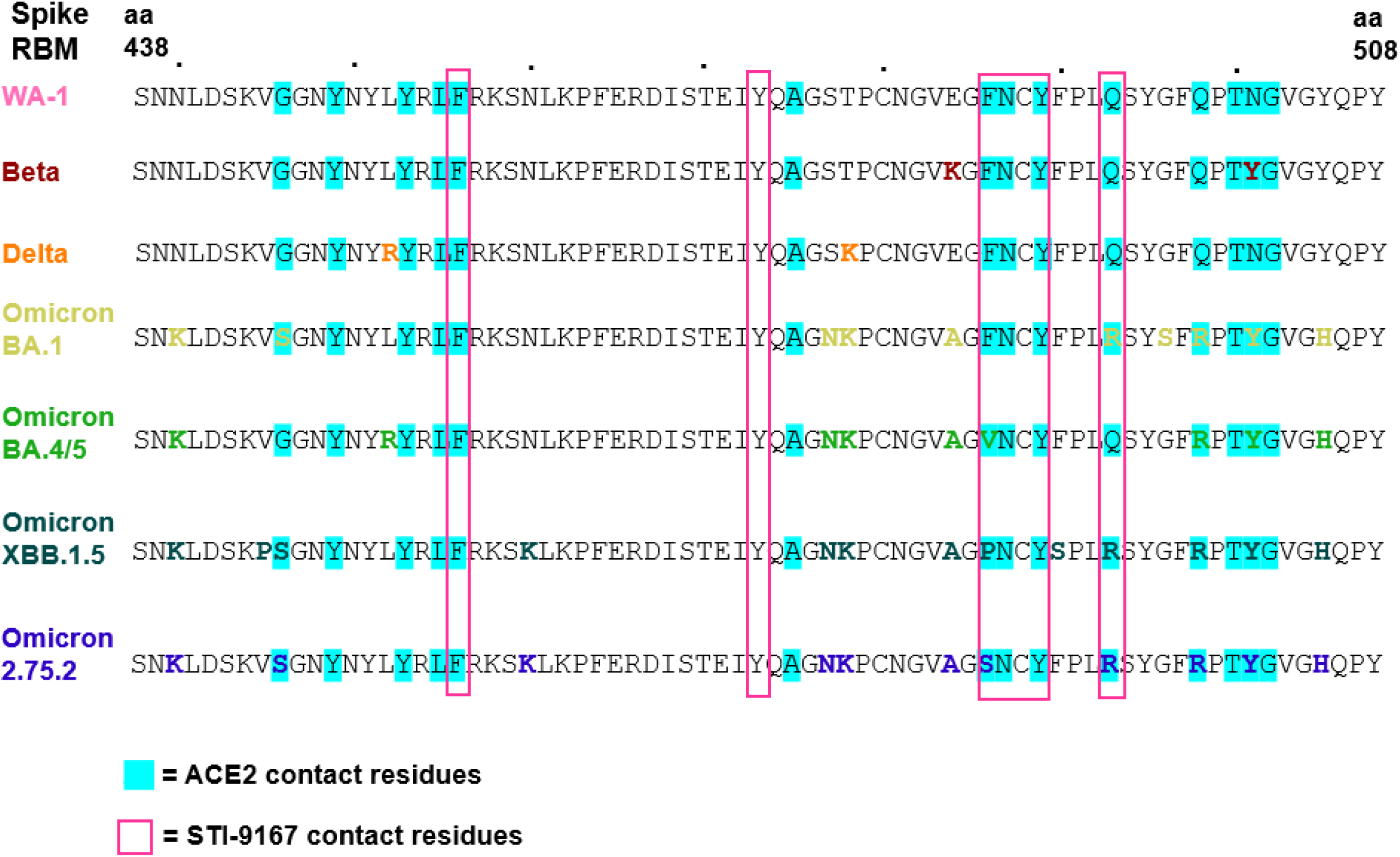
Comparison of the STI-9167 Fab contact residues with the Spike receptor-binding motif (RBM). The ACE2 (cyan highlights) and STI-9167 (pink boxes) contact residues of the Spike receptor-binding motif (RBM, aa 438-508) of SARS-CoV-2 WA-1, Beta, Delta, Omicron BA.1, Omicron BA.4/5, Omicron XBB.1.5, and Omicron 2.75.2 are shown. The amino acid sequence of the RBD is represented using the single letter amino acid code.

Knowledge of the nmAb’s epitope and its molecular underpinnings allows us to preemptively predict the location and nature of mutational variants in Spike that would abolish antibody neutralization, as well as those which the antibody could likely tolerate. This more broadly indicates the value of experimentally soliciting viral escape mutants to neutralizing antibodies, not only for determining those nmAbs’ epitopes, but also as a means for predicting potential mutations that may arise as SARS-CoV-2 variants continue to proliferate under dynamically-changing conditions of evolutionary fitness. Based on the cryo-EM map alone, we might surmise that a mutation at position 486 to a polar residue would abrogate neutralization by FG-10A3/STI-9167, whereas a different hydrophobic substitution at the same position may still be recognized; indeed, neutralization assays to determine FG-10A3’s activity against rcVSV-S containing point mutations at position 486 demonstrated that, while introduction of the polar residue serine completely impaired antibody neutralization, substitution with nonpolar residues such as leucine and valine still permitted FG-10A3 recognition and viral neutralization (**Figure 4**).

Strikingly, despite the major differences in their neutralization profiles and recognition of distinct epitopes within the Spike RBD, these nmAbs were all generated using the same immunization strategy, that of inoculating transgenic humanized mice with SARS-CoV-2(WA-1) Spike RBD protein. Discovery and production of human nmAbs in transgenic animals poses several distinct advantages, such as permitting *in vivo* affinity maturation and clonal selection for subsequent antibody optimization (40). This work highlights the use of transgenic mice and the strategy of immunization with a protein domain as a means of generating a diverse panel of nmAbs that target a specific antigen, as well as illustrating the potential for such nmAbs to complement each other’s neutralization capabilities in order to most effectively prevent infection and spread of viral variants.

## Acknowledgments

Some of this work was performed at the National Center for CryoEM Access and Training (NCCAT) and the Simons Electron Microscopy Center located at the New York Structural Biology Center, supported by the NIH Common Fund Transformative High Resolution Cryo-Electron Microscopy program (U24 GM129539), and by grants from the Simons Foundation (SF349247) and NY State Assembly.

## Author contributions

K.E.A. designed and performed the neutralization and antibody resistance studeis. L.B. and S.K. designed and generated rcVSV with SARS-CoV-2 spike variants. C.S.S and C-T.H. designed and performed the sequencing analysis of antibody-resistant rcVSV strains. Y.F, R.L., L.T., and R.A. designed and generated the cleavage and purification of the Fab fragment used in cryo-EM. G.B. designed the cryo-EM experiments. J.A.D designed and performed the antibody binding studies. D.C.A. and G.B. performed cryo-EM experiments. G.B. analyzed cryo-EM data. K.E.A., L.B., C.S.S., G.B., R.A., J.A.D, and B.L., and D.T. wrote and edited the manuscript. D.T. conceived and supervised the project.

## Materials & Correspondence

Requests for material should be addressed to Domenico Tortorella and Goran Bajic. The EM maps have been deposited in the Electron Microscopy Data Bank (EMDB) under accession code EMD-28537 and the accompanying atomic coordinates in the Protein Data Bank (PDB) under accession code 8EQF. The aligned micrographs are available on the Electron Microscopy Public Image Archive (EMPIAR) under accession number EMPIAR-11341.

